# Food Microbiome Metabolic Modules (F3M), a tool suite for functional profiling of food microbiomes

**DOI:** 10.1101/2025.07.31.667869

**Authors:** Julien Tap, Nacer Mohellibi, Colin Tinsley, Valentin Loux, Stéphane Chaillou

## Abstract

The study of microbial metabolic interactions within food microbiomes represents a key scientific approach for improving the quality and health benefits of food. In such studies, methods based on gene expression levels (metatranscriptomic) analysis are promising. However specific tools are required to overcome the challenges posed by food microbiomes, in particular the high variability of microbiomes between samples and the difficulty of automatically inferring the annotation of metabolic functions across taxa. To adress this gap, we present the Food Microbiome Metabolic Modules (F3M) tool suite, which comprises (1) a curated database containing about 1,985 functional genes representing key fermentative metabolic reactions in food microbiomes, (2) a F3M Builder for generating F3M-annotated gene catalogs and mapping of metatranscriptomic reads, and then (3) an F3M R package to parse and aggregate gene expression data by taxonomic and functional categories for downstream analysis. The F3M taxonomy is organized according to the Genome Taxonomy Database (GTDB) nomenclature, whereas the F3M functional repertoire is structured hierarchically into 183 metabolic modules, which enable multi-scale analysis of inter-organism metabolic interactions and meaningful fermentative outputs (e.g., primary alcohols, acetate). Notably, a dedicated ‘redox’ module captures oxido-reduction mechanisms and NADH-dependent pathways central to fermentation, while an ‘uptake’ module complements the metabolic pathways to trace potential metabolite exchanges across taxa. Together, the F3M suite provides a robust framework for uncovering functional dynamics within food microbiomes. The F3M tool suite is available as open-source.

## Introduction

There are more than a thousand different fermented foods in the world (Gänzle, 2022), made from a wide variety of different products (vegetables, cereals, meat, dairy products, fruit, *etc*.) and obtained from numerous recipes that are, in most cases, reflections of an ancestral and regional culture. Each fermented food results from a microbial ecology process that generates a well-defined type of microbial community. Characterizing these communities is essential to understanding their dynamics, as well as providing opportunities to improve safety and sustainability, and in this context, the development of databases such as FoodMicrobionet (Parente et al., 2019) or the Food Microbiome Database (Carlino et al., 2024), has allowed a better assessment of the broad food-borne microbial diversity.

However, understanding the interactive metabolic nature of these microbial food communities remains a challenge in its own right and one that has yet to be fully resolved. Among the reasons for this difficulty is that foods are open systems and are dynamic, consisting of successive populations. Unlike other microbiomes, the best studied being the gut microbiome, whose species structure reflects co-evolution with the host, food microbiomes are far more variable in nature.

In the case of food microbiomes, functional inference from the metagenomic data can lead to a bias in the interpretation of community functioning, as the functional capacities carried by certain species may reflect activities linked rather to their life-cycle in their environments of origin (*e*.*g*., raw materials, processing plants) than to their function in the foodstuffs. Moreover, many food studies focus on the end product (fermented or spoiled), and neglect the dynamics of population succession. This leads to major gaps in our knowledge of metabolic interactions between the microbial strains involved throughout the food fermentation process.

Metatranscriptomics is probably the most appropriate method for studying metabolic interactions within food microbiomes. It provides a more focused view of the functions expressed by the community at different stages of the dynamics. However, the number of available food metatranscriptomic datasets is still very small. While it is possible to list more than 2,500 food metagenomic samples (Carlino et al., 2024), only 27 food metatranscriptomic samples are listed in public data archives such as MGnify (Richardson et al., 2023) when writing this article.

Several methodological constraints complicate the use of metatranscriptomics in food products. Recovering good-quality microbial mRNA from solid-state food material is certainly one of the most important. This difficulty is compounded by frequent contamination with RNA originating from the food matrix; extraction techniques must be reassessed depending on the matrices which are to be investigated. Another important constraint is the difficulty of exploiting data at both functional and statistical levels. Indeed, the high variability in the composition of microbial communities between biological replicates (different samples) of a given food can generate difficulties in the comparative statistical analysis of gene expression levels. For example, statistical tools widely used for this type of analysis, such as DESeq2 (Love et al., 2014), use very specific normalization procedures that take into account the broad dynamics of gene expression levels (up to 10^6^-fold), the library size and the gene-wise dispersion to estimate the scaling factors for each sample. In this type of analysis, the number of biological replicates is crucial for statistical validation. However, as shown in Figure 1A, if the variability of the community is too great between replicates, this can lead to a significant change in the number of genes to be normalized between replicates and, consequently, produce a zero-inflated matrix that the statistical models have difficulty in handling, hence leading to biased or skewed normalized data. Thus, if differences in species abundance between food microbiomes replicates becomes too great, it will usually disrupt and make any normalization step ineffective between the different conditions analysed. One solution to this problem is to consider functions or pathways rather than genes, as illustrated in Figure 1B. Since some metabolic functions are shared between several species, each contributing collectively to the required activity, aggregating the read counts to the scale of a function minimizesthe impact of variation in species abundance on data normalization. This approach requires that functional annotations of genes be of high quality and identical across distant taxa.

**Figure 1.**
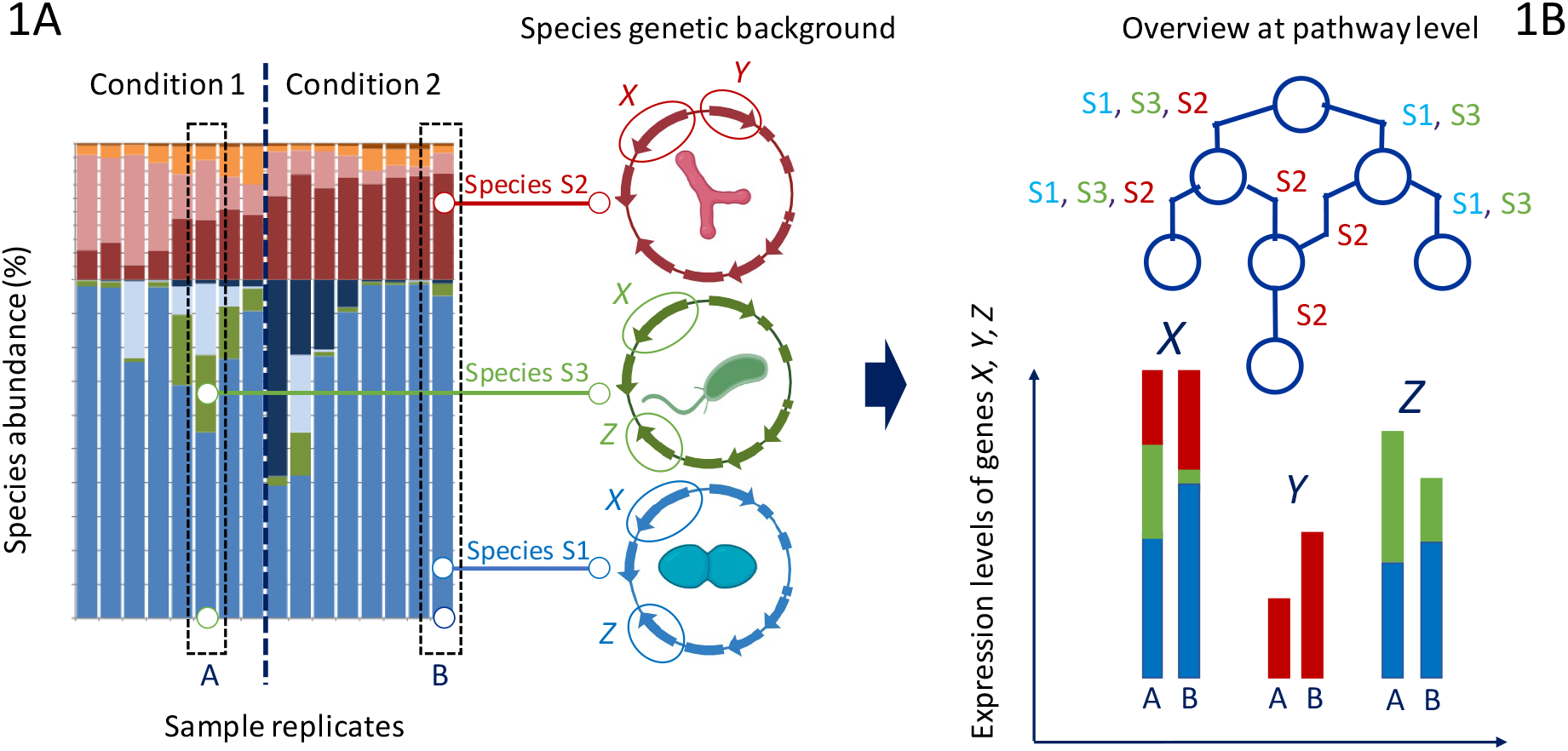
Overview of the challenge of studying food microbiomes at the functional level from gene expression data. Figure 1A illustrates how the variation of species abundance in fermented food replicates can be very high. The example shows two conditions being compared with eight replicates performed for each. The relative abundance of various species is depicted by colors. As each microbiome species also carries a specific genetic background for metabolism, combining both features often skews the statistical comparison between two experimental conditions because functional genes *X, Y*, and *Z* are not shared uniformly between the species. Figure 1B illustrates how an overview of the metabolic pathway could help to overcome these biases (for simplicity, sample A from condition 1 is compared to sample B from condition 2). At the community level, the expression of gene X (representing one branch of the pathway shared by the three species) might not be differentially expressed at the level of the community despite species abundance variability, whereas genes Y and Z (representing different representatitivity among the species for other branches of the pathway) might show significant differences in expression at the level of the community.

Carrying out metatranscriptomic studies in this context, particularly when working on ecosystems that have not been defined *a priori*, requires generic tools that enable a global functional analysis of databases derived from metagenomes and metatranscriptomes.

An example of such a tool is the ChocoPhlan3 pipeline integrated into the HumanN 3.0 suite, which incorporates taxonomic (species-level) information and metabolic function (Beghini et al., 2021). HumanN 3.0 uses native UniRef90 annotations from ChocoPhlan species pangenomes corresponding to 549 MetaCyc pathways and 2895 EC numbers. Another pipeline, Super-Focus (Silva et al., 2016), aligns metatranscriptomic input reads against a subset of the SEED database (Overbeek et al., 2005) based on functional subsystems. Sometimes, these tools are integrated into dedicated metatranscriptomic workflows such as metaPro (Taj et al., 2023) or SqueezeMeta (Tamames and Puente-Sánchez, 2018). Among these tools, HumanN 3.0 is the only one that, thanks to the aggregation of read counts at both taxonomic and functional levels (two levels including the level of MetaCyc pathways and the level of Uniref90 gene families, each gene family being potentially assigned to a metabolic reaction in KEGG and/or MetaCyc), offers an analytical strategy capable of providing a partial answer to the problem presented in figure 1.

What all these tools have in common is to use as a reference the Kyoto Encyclopedia of Genes and Genomes (KEGG) (Kanehisa et al., 2023) and MetaCyc (Caspi et al., 2020) databases, which are the most widely used in metabolic studies. KEGG and MetaCyc have the advantage of being generic and extensive databases. However, they have the disadvantage of being focused on metabolic pathways based on humans or on model or pathogenic organisms that are well documented in the literature. Another drawback is that the assembly of functional categories or metabolic pathways is not always the same between the two databases. In addition, many metabolic pathways and functions are poorly identified in these databases, particularly those pertaining to anaerobic and/or fermentative metabolisms. The knowledge gap in this area has recently been clearly highlighted (Hackmann and Zang, 2023). Thus, automatic functional inference with overly generic tools is far from optimal when considering fermentative microbiome datasets.

This problem is not new and has been the subject of the creation of dedicated tools such as human-gut metabolic modules (Vieira-Silva et al., 2016), the gut-brain modules (Valles-Colomer et al., 2019), and, more recently, the enteropathways (Shiroma et al., 2024). These modules, some of which are now widely used for the functional study of human-associated microbiomes, have been built using strategies that make use of KEGG orthology (KO) sets, which represent the most widely used and informed metabolic annotations, together with expert curation of functions based on literature analyses. These functions are then organized into metabolic modules or pathways, which are, in some cases, similar to those proposed by KEGG or MetaCyc, and, in others, organized differently to allow monitoring of functional variations from an angle focused on the digestive functions of the gut microbiota.

In contrast, there are currently no similar modules for food microbiomes, even though the microbial ecology of foods also has its specificities. In particular, food environments are predominantly colonized by lactic acid bacteria and/or yeasts, whose metabolic functions in fermentation are poorly annotated or even incorrectly positioned in the KEGG or MetaCyc modules.

Here we present a new tool, F3M, created with a triple objective: firstly, to perform expert curation of functional annotations related to the fermentative metabolism of food bacteria; secondly, to aggregate these functions into new modules providing comprehensive coverage of pathways in food microbiomes and thirdly, to develop a suite of tools allowing more straightforward and more relevant statistical analysis of food microbiome metatranscriptomics data which can be performed at several levels of grouping of functional and/or taxonomic data. With this last objective in mind, we wanted to create an open-source tool suite that, rather than being very generic, can be user-adapted and customized to the ecosystem under study.

## Material and methods

### Development of the F3M tool suite

As described in Figure 2, the F3M tools suite comprises one specific knowledge database and two additional computational methods and data managers: the “F3M builder” on one hand and the “f3mr” R package on the other hand. The F3M suite is open-access and was constructed to allow easy use and development by users. The F3M builder was developed (1) to build user-defined gene catalogs with F3M functional annotations, which may be specific to a given food microbiome, and then (2) to allow easy mapping of metatranscriptome sequence reads onto these. Then, data from the mapping is processed using the f3mr R package for downstream analysis.

**Figure 2.**
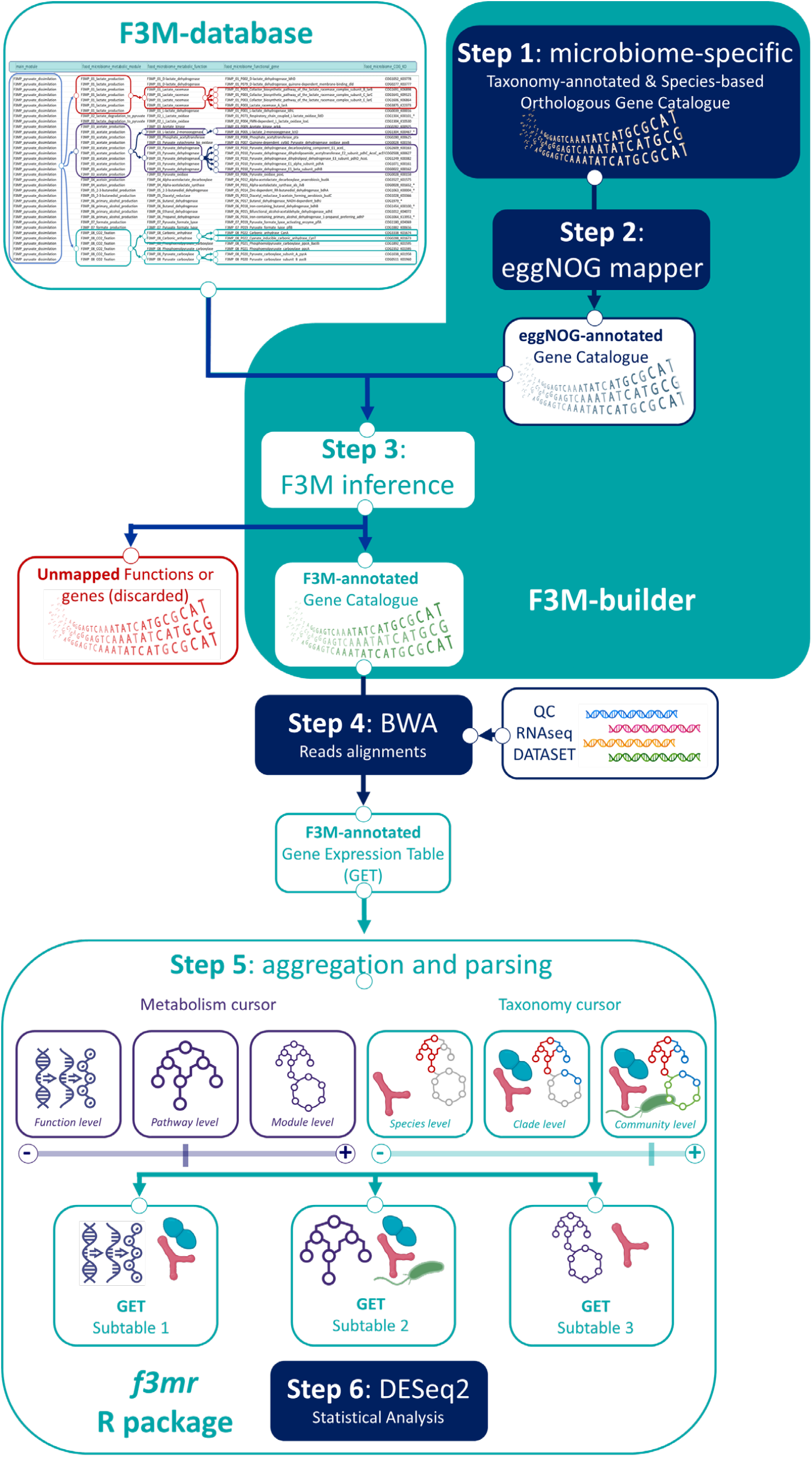
Overview of the F3M tool suite with the three main elements (database, builder, and R package) and the six main steps of the workflow. Steps requiring dependencies to other software or databases such as ProGenome (Mende et al., 2017), eggNOGmapper (Cantalapiedra et al., 2021), bwa (Li and Durbin, 2009), DESeq2 (Varet et al., 2016) are indicated in dark blue.

### Curation of functional annotations and creation of the F3M database

A large collection of bacterial pangenomes extracted from the proGenome V3 database (Mende et al., 2017) and belonging to food-associated bacteria (about 80 different bacterial families) was selected to curate the functional annotation. This collection was chosen based on the current knowledge of food microbiomes available in FoodMicrobionet (Parente et al., 2019) or the Food Microbiome Database (Carlino et al., 2024). The curation was performed in three steps: first, the pangenomes were submitted for automatic annotation by eggNOG-mapper tools (Cantalapiedra et al., 2021). Secondly, the process of curation comparing eggNOG-mapper KEGG KO and Eggnogg COG identifiers consisted in clustering these COG_KO combinations across the taxonomic distances of the pangenomes, and thirdly, we linked these combinations to genomic contexts and functional data present in the scientific literature in order to determine the functional annotation correctly and associated this annotation to the COG_KO identifier. The F3M database and the metabolic maps depicting the functional modules are freely available (see data, script and code availability section).

### Development of the F3M builder

The F3M builder is a set of Snakemake workflows and Python scripts divided into two specific parts: (1) the catalog construction workflow, and (2) the reads mapping workflow. Complete documentation concerning the steps of the F3M builders, data inputs, outputs, tool dependencies and configuration, is available as open-source (see data, script, and code availability section).

The first workflow is called “*build_a_gene_catalog_from_pangenomes*”. It is used to build a food microbiome-specific gene catalog with annotations that represent the F3M modules, functions, and genes (see result section). To this end, it first downloads the pangenomes of the bacterial species known to populate a specific food microbiome from the proGenome v3 database (Mende et al., 2017), this list being provided by the user. The proGenome V3 database allows efficient parsing of pangenomic data associated with accurate, curated clusters of genomes per species, which are defined in terms of a so-called specI number (Mende et al., 2013) together with a complete taxonomic profile. Subsequently, the orthologous genes are clustered at the species (specI) level to produce a species pangenome, which is then annotated using the eggnog mapper resources (Cantalapiedra et al., 2021) and information from the F3M database. Two examples of open-access F3M gene catalogs are provided (see data, script, and code availability section), one for meat microbiomes and the second for fermented vegetable microbiomes.

The second workflow, called “*map_reads_to_gene_catalog*” is used to quantify, from a metatranscriptomic dataset, the level of gene expression corresponding to these F3M-curated annotations and to assign the species involved in these functions. Therefore, the F3M builder has three main outputs: two files corresponding to the annotations of the F3M gene catalog (one for taxonomy and another for metabolic functions) and the mapping files. These files serve as input to the f3mr R package.

### Development of the f3mr R package

The f3mr package can be executed in a controlled environment using R (version 4.4) with Tidyverse tools (version 2.0) (Wickham et al., 2019) and the Targets package (version 1.7) (Landau et al., 2021) for workflow management. See data, script and code availability section for the complete documentation and installation guide.The package is roughly divided into four steps:

- The first step comprises the “build_ref_db()” function to construct a reference database from functional annotations and taxonomic classifications obtained from the F3M builder, “build_a_genes_catalog_from_pangenomes*”* workflow.
- The second step comprises the import of the read counts obtained from the F3M builder, “map_reads_to_gene_catalog*”* workflow. Two functions can import the data, either the import_sample_count() or the import_multiple_samples().
- The third step is the aggregate_counts() function, aggregating counts at taxonomic and functional levels chosen by the user.
- The fourth step is the build_count_matrix() function, which creates a count matrix suitable for downstream analyses such as DESeq2 (Varet et al., 2016).

The f3mr package offers a tutorial (see complete documentation) based on a shallow-depth (∼ 400K reads) metatranscriptomic dataset from beef carpaccio mapped against the meat_microbiomes_F3M_genes_catalog_V1.1.0 in order to allow the user a rapid familiarization with the tool.

### Analysis of the beef carpaccio dataset

The link to the full description of the dataset can be found at the data, script and code availability section. The data were processed as follows: quality filtering of the fastQ RNA-seq reads was performed with fastp (version 0.24.0) (Shifu et al., 2018) using default options. Filtered reads from the eight samples were then assembled *de novo* with Spades (version 4.1.0) (Nurk et al., 2017) using the *meta* option. Gene detection was then performed on the assembled contigs with Prodigal (version 2.6.3). These *de novo* assembled genes from the dataset were used as a reference for mapping the filtered-fastQ reads using bowtie2 (version 2.5.4) (Langmead et al., 2012). The quantity of reads mapping to the whole set of microbial genes was estimated and was compared to the results obtained when using F3M or HumanN 3.0 (version 3.9) function (Beghini et al., 2021) mapping; the latter was used with the standard options. Differential analysis was performed with the DESeq2 package (Varet et al., 2016). Raw outputs from the F3M suite, HumanN 3.0 mapping, and subsequent DESeq2 analysis on the beef carpaccio dataset can be found at the data, script and code availability section.

## Results

### Rationale and general architecture of the Food Microbiomes Metabolic Modules

The F3M is based on three elements, which are illustrated in Figure 2. Firstly, a repertoire of metabolic functions whose annotations have been expertly curated is organized hierarchically, with a classification that evolves from the broad main functional category through functional module (metabolic module or pathway), to metabolic function and functional gene. This organization is leads to a codified nomenclature and constitutes the F3M database.

The second element is based on one or more catalogs of orthologous, non-redundant genes at the pangenome scale of a given species, which are annotated taxonomically and on which the F3M annotations will be transferred. These F3M gene catalogs represent the nucleotide references on which metatranscriptome reads will be mapped to obtain a table of gene expression levels. In this matter, we have built the F3M suite to be highly flexible and adaptable to the food microbiome studied (Figure 2).

Several strategies can be adopted for the creation of a gene catalog. A very inclusive approach is to create a gene catalog covering the known diversity of genes and microbial species that have been characterized in foods. For example, this catalog could be created from the over 800 species-level genome bins (SGBs) characterized in the recently published Food Microbiome Database (Carlino et al., 2024). However, the larger the catalog, the greater the computational resources required for read mapping. An alternative strategy, which we propose here, is to create catalogs dedicated to certain food ecosystems on which a given study focuses. With this second approach in mind, we present two examples of gene catalogs annotated by F3M. One is dedicated to the study of spoilage microbiota in meat products, the other to the microbiota of fermented vegetables such as sauerkraut or kimchi. To build these custom catalogs, we extracted a list of key bacterial species identified in these microbiomes (see materials and methods section) and we built up the catalogs from species’ pangenomes extracted from the proGenome V3.0 public database (see data availability section for the link to further description of these two catalogs).

The third element of the F3M strategy is based on the concept of aggregating the mapping data according to two parameters: one metabolic and functional, the other taxonomic and structural (Figure 3). This aggregation is accomplished with the help of a parser (f3mr R package) that uses the codified nomenclature of the F3M database and the GTDB taxonomy (Parks et al., 2022) to produce different Gene Expression Tables (GET), which in turn serve as input for differential statistical analyses. This dual-cursor concept is one of F3M’s assets for studying the various levels at which metabolic interactions or functional changes within food microbiomes take place.

**Figure 3.**
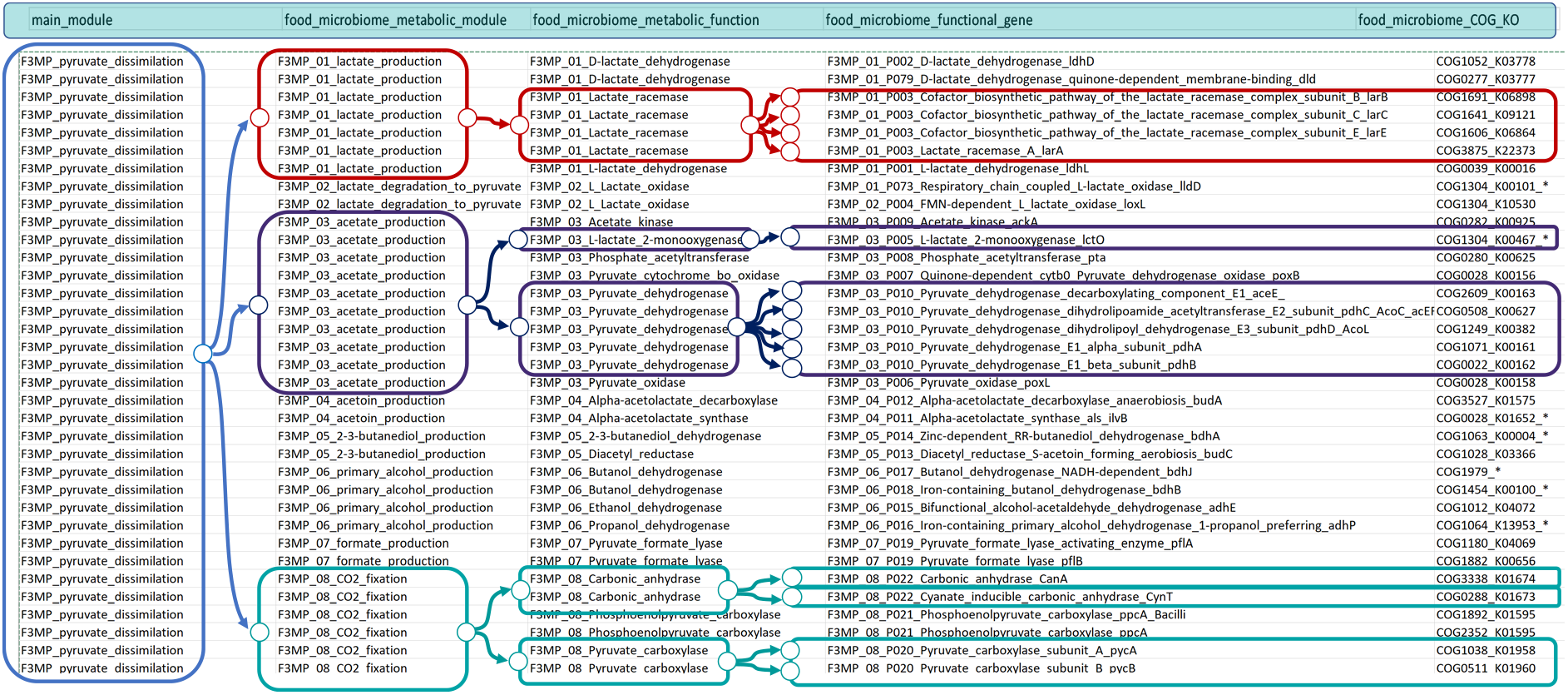
Overview of F3M database and its hierarchical nomenclature. Several examples are shown with colored boxes starting from the broadest functional grouping on the left (main_module), here illustrated by the F3MP_pyruvate_dissimilation module, to more specific ones (food_microbiome_metabolic_functional_gene) at the right. In the case of F3MP_01_lactate racemase (uppermost, red boxes), the figure shows that the metabolic function is represented by several genes (multimeric complex). This hierarchical structure of the F3M database allows the aggregation of expression data from the four genes into a single function. The last column of the database indicates the COG_KO identifiers used to infer the functions.

### Strategy for curation of functional annotation and the establishment of the F3M database codified nomenclature

The curation of metabolic functions in F3M is based on a combination of COG identifiers and, as with gut metabolic modules already published (Vieira-Silva et al., 2016), one or more KO identifiers from the KEGG database. This combination is easily obtained by automatic annotation tools such as eggNOG mapper. The next step is to annotate this COG_KO combination with reference to different databases such as MetaCyc or KEGG, superimposing an expert approach based on genetic context analysis and bibliographic data.

The important point in this curation step of the F3M database is that each function in the database is associated with a unique COG_KO combination.

To illustrate our approach, we may use a few examples chosen from those shown in Figure 2. For instance, we have annotated two functional genes carrying the carbonic anhydrase function (both coded under the food_microbiome_functional_gene designation F3M_08_P022). These two genes have distinct COG-KO combinations (COG3338_K01674 and COG0288_K01673). This approach makes it possible to distinguish two classes of bacterial carbonic anhydrases: generic (encoded by the canA gene) and cyanate-inducible (*cynT*). In the F3M nomenclature, the expression of these two functional genes can be aggregated under the same food_microbiome_metabolic_function, F3MP_08_carbonic_anhydrase.

The other example illustrates the annotation of the various lactate oxidases, encoded by the COG and KO identifier combinations COG1304_K10530, COG1304_K00101_*, or COG1304_K00467_*. The use of the wild card (*) for the COG1304_K00101_* combination, for example, indicates that the automatic annotation may associate other KOs with K00101 (notably K10530), but that only K00101 is specific to the *lldD* gene encoding the lactate oxidase associated with the aerobic respiratory chain. Codification COG1304_K10530 (K10530 strictly selected) has been appraised as the FMN-dependent lactate oxidase *loxL* of lactic acid bacteria. Nevertheless, these two functional genes can be aggregated under the same function (F3MP_02_lactate_oxidase) as they are both involved in the oxidation of lactate to pyruvate (module F3MP_02_lactate_degradation_to_pyruvate). In contrast, the third functional gene (L-lactate_2_monooxygenase_lctO) is associated with a different functional module (F3MP_03_acetate_production). This use of the wild card (*) facilitated the extraction and sorting of automatic annotations to better associate them with expert functional information and, *in fine*, functional curation.

### Construction of the F3M modules in the database

The aim of redesigning the functional modules (metabolic pathways or functional networks) compared to those proposed by existing databases such as MetaCyc or KEGG was to create modules that were more relevant to understanding microbial interactions, specifically within foods, with a focus on fermentation, a predominantly anaerobic process. For instance, we have created a main module linked to pyruvate dissimilation, a key metabolic node in fermentations. Although a similar module exists in MetaCyc (mixed acid fermentation) or KEGG (pyruvate metabolism), the revised F3M version we propose for this module integrates a more complete view of associated pathways, for example, methylglyoxal reduction, as well as malolactic and citrolactic fermentation pathways. It also integrates an exhaustive list of dehydrogenases involved in producing primary alcohols (ethanol, propanol, butanol, etc.).

In addition to the strategy of redesigning functional modules in the light of food microbiome specificities, we also created two new modules that we see as essential to studying metabolic interactions. As illustrated in Figure 4, redox balancing of NAD+/NADH, or redox reactions in general, is a central and essential processes that govern microbial metabolism. Although many functions are well described in both MetaCyc and KEGG databases, they are not connected comprehensively. We have therefore created a redox-focused module (see Figure 5) which connects about one hundred functional genes involved in different redox processes and enables a better understanding of these activities, which Bouranis and Tfaily have recently classified as the microbial black box (Bouranis and Tfaily, 2024).

**Figure 4.**
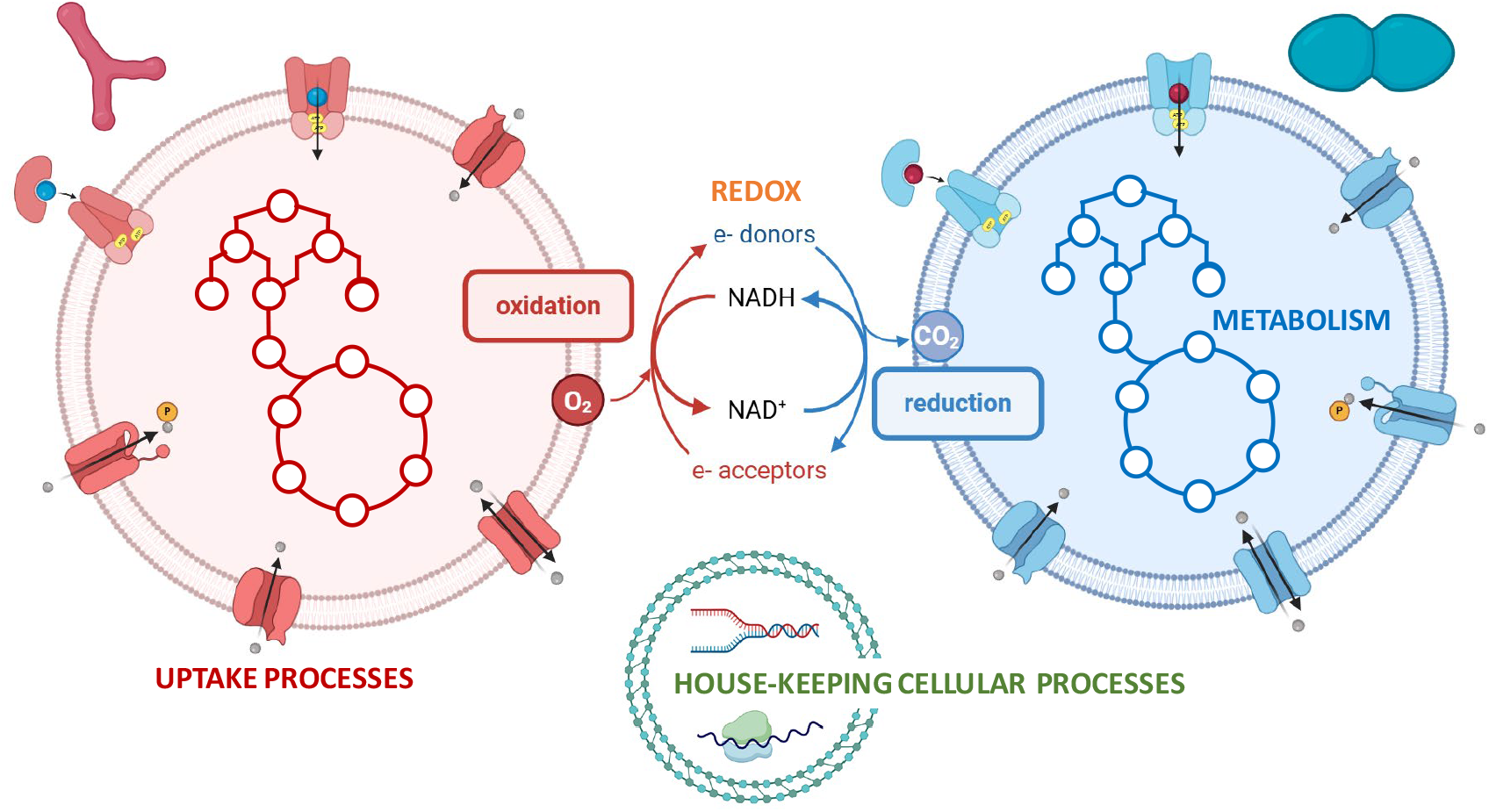
Overview of different classes among the F3M main functional modules allowing the linkage of the major functional processes in microbial interactions: the modules for metabolism (illustrated by the pathways within the two bacterial cells), redox processes (central oxido-reduction mechanisms between the two cells), and uptake processes (various transporters in the cell membrane of the two cells). Housekeeping cellular processes modules are also provided in the F3M but are dedicated to the normalisation steps commonly performed in metatranscriptomic statistical analysis.

**Figure 5.**
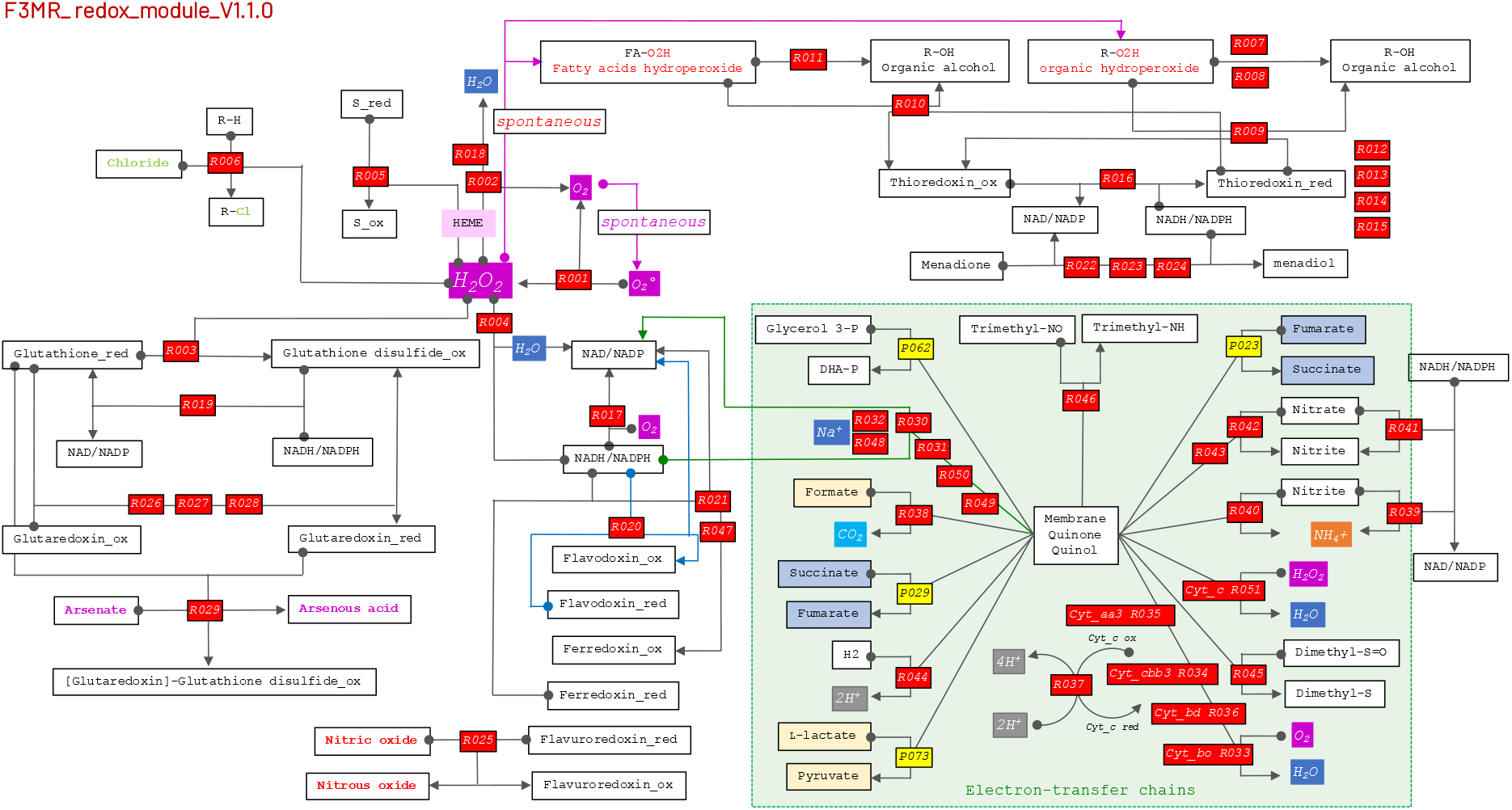
Example of an F3M metabolic map. Such maps are available for each module excepting those dedicated to house-keeping genes. The maps are not yet an output of the FM3 tool suite, but they are provided as supplementary documents at the F3M database repository (see data availability section) to help F3M users visualize the structure of the various metabolic modules and the corresponding metabolic reactions in the modules. This example shows the redox module, whose functions are organized peripherally around the NAD/NADH central component. The food_microbiome_functions of this module are depicted in red boxes and by their code in the F3M database (for instance, R002 stands for F3MR_01_R002_Heme-dependent_catalase_katA) involved in peroxide metabolism. Electron-transfer chains are grouped in the green box with dehydrogenase components on the left hand side and reductase (oxidase) components on the right. In some cases, metabolic reactions may belong to several modules. In these cases, we have assigned the functions to only one primary module, although for the purposes of visualisation we retain their association with each of the two or more modules, meanwhile keeping track with the boxes’ color-code. Such an example can be found here, with P073 (yellow color) being the respiratory-chain-dependent L-lactate oxidase lldD, which belongs primarily to the pyruvate dissimilation module.

We also created a module that covers all the processes involved in metabolite uptake. These processes are also key to exchanges between micro-organisms. The MetaCyc database provides little information on these processes, while the KEGG database, in contrast, lists numerous processes organized by orthologous gene families. Uptake-associated functional genes constitute a complex set of gene families, sometimes highly conserved, whose functional assignment may be challenging. We therefore adopted an intermediate strategy, creating a panel of 20 sub-modules that allow specific classification of major transport systems by metabolite category. More than 100 uptake functions (> 400 functional genes) have been assigned to these modules, including a category rarely found in metabolic databases and linked to the Energy-Coupling-Factor-type ATP-binding transport systems, whose functional importance for the transport of vitamins or mineral co-factors is now well established (Rempel et al., 2019).

Finally, a set of main modules related to cellular housekeeping processes (replication, translation, *etc*.) has been built to create a fairly large corpus of genes conserved between distant taxa and involved in cellular housekeeping functions. The primary role of this set of functions is to allow normalization of gene expression levels. A summary of the F3M database modules is given in Table 1 below.

**Table 1.** Summary of the F3M database, from food_microbiome_main_module to food_microbiome_functional_gene.

| main_module | metabolic_module | metabolic_function | functional_gene | Type |
| --- | --- | --- | --- | --- |
| F3MB_nucleotide_metabolism | 08 | 59 | 73 | metabolism |
| F3MC_carbohydrate_metabolism | 29 | 169 | 201 | metabolism |
| F3MF_cofactors_metabolism | 16 | 144 | 164 | metabolism |
| F3ML_fatty_acids_metabolism | 05 | 25 | 36 | metabolism |
| F3MN_amino_acid_nitrogen_metabolism | 35 | 224 | 267 | metabolism |
| F3MP_pyruvate_dissimilation | 21 | 84 | 138 | metabolism |
| F3MR_redox | 11 | 47 | 118 | redox |
| F3MU_uptake_transport_systems | 20 | 113 | 430 | uptake |
| Σ metabolism+redox+uptake = 8 modules | **145** | **865** | **1,427** |  |
| F3MD_DNA_metabolism | 07 | 29 | 129 | House-keeping |
| F3ME_ATP_biosynthesis | 02 | 03 | 16 | House-keeping |
| F3MM_cell_motility | 03 | 29 | 48 | House-keeping |
| F3MA_RNA_metabolism | 04 | 18 | 97 | House-keeping |
| F3MS_cell_division_and_shape | 05 | 21 | 47 | House-keeping |
| F3MT_translation_and_posttranslation_processes | 13 | 52 | 170 | House-keeping |
| F3MW_cell_wall_metabolism | 04 | 35 | 51 | House-keeping |
| Σ house-keeping processes = 7 modules | **38** | **187** | **558** |  |
| Σ all modules = 13 modules | **183** | **1,052** | **1,985** |  |

Note: the numbers refer to the number of entries in each category

### Beef carpaccio food metatranscriptomic dataset as an F3M tool suite tutorial

A dataset is available for rapid testing of the F3M tool suite and as a tutorial. A detailed description of this dataset is available (see data, script, and code availability sections). It corresponds to eight metatranscriptomic samples from a beef carpaccio microbiome stored for 4 days at 4°C under two storage conditions (air or vacuum, four replicates for each condition).

Briefly, this dataset allows the analysis of potential metabolic interactions between the two dominant species in these microbiomes and highlights the added value of the various F3M modules to these studies. We also analyzed this dataset with HumanN 3.0, the tool most similar to F3M in its analytical strategy, in order to compare the outputs of the two methods. The dataset represents a simplified ecosystem (Table 2) in which the two main species have different sensitivities to the presence or absence of oxygen: *Latilactobacillus sakei* (fermentative species) and *Pseudomonas fragi* (aerobic species).

**Table 2.**
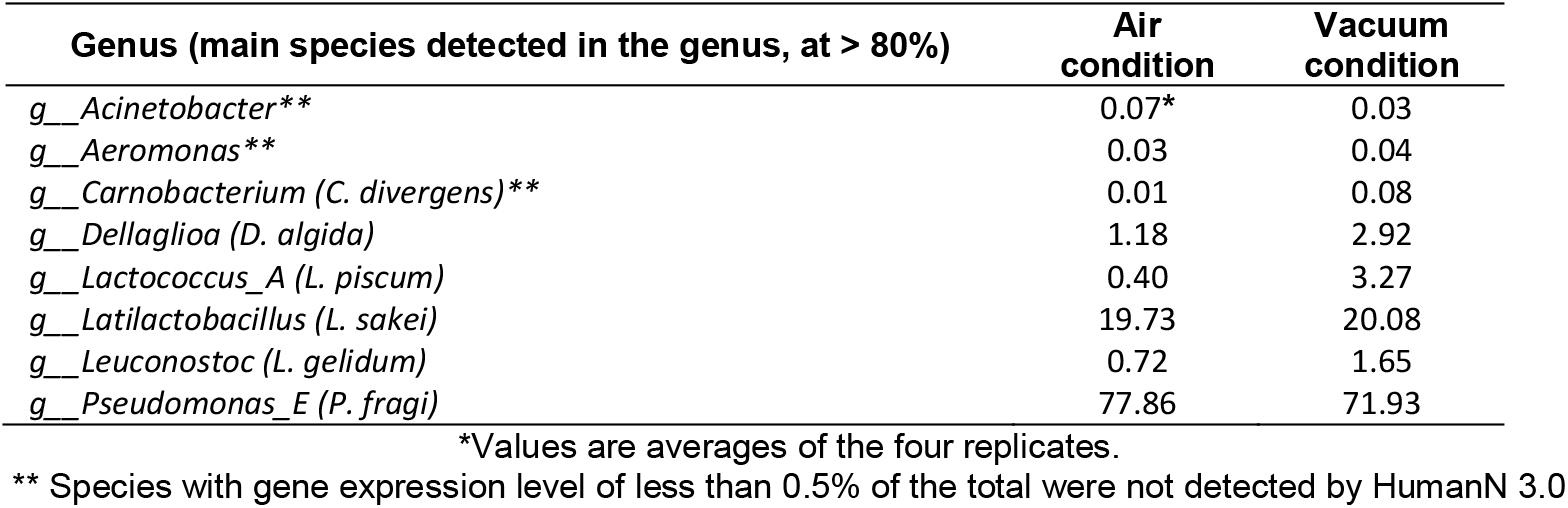
Relative levels of gene expression within the microbiota aggregated at the genus level within the beef carpaccio metatranscriptomic test dataset.

The results in Table 2 show that despite the different storage conditions, the two species have their average gene expression at almost identical levels between the two conditions, *i*.*e*., around 20% of total reads for *L. sakei* and 70-78% for *P. fragi*. The observation that *P. fragi* remains highly competitive in vacuum storage was not expected based on the different sensitivity of the two species to the presence or absence of oxygen. Therefore, we used the F3M tool suite to test the hypothesis of a metabolic interaction that could benefit *P. fragi* in the vacuum storage condition. To compare HumanN 3.0 and F3M methods we focused our analysis on two levels which represented two similar levels of aggregation between the two methods. For both F3M and HumanN 3.0, we compared the taxonomy at genus versus community level, and for the functional comparison we chose two additionnal levels which were: functional genes (F3M)/Uniref90 GeneFamilies (HumanN 3.0) versus F3M modules (F3M)/pathways (HumanN 3.0). Table 3 shows an overview of the output resulting from the mapping at these different levels.

**Table 3.** Comparative overview between HumanN 3.0 and F3M tool suite on the mapping results of the beef carpaccio metatranscriptomic dataset.

| Percentage of mapping reads | Air condition |  | Vacuum condition |
| --- | --- | --- | --- |
| F3M - Overall percentage of mapping reads** | 59.6 ± 5.4* |  | 58.8 ± 7.3 |
| HumanN 3.0 - Overall percentage of mapping reads** | 80.6 ± 1.3 |  | 68.9 ± 11.3 |
| Overall percentage of *L. sakei* reads mapping to F3M functional genes § | 55.9 ± 1.8 |  | 50.5 ± 4.7 |
| Overall percentage of *L. sakei* reads mapping to HumanN 3.0 gene families § | 32.6 ± 1.3 |  | 31.8 ± 3.4 |
| Overall percentage of *P. fragi* reads mapping to F3M functional genes § | 58.9 ± 1.8 |  | 54.7 ± 2.9 |
| Overall percentage of *P. fragi* reads mapping to HumanN 3.0 gene families § | 90.8 ± 2.8 |  | 73.1 ± 10.0 |
| Number of different functional categories identified by mapping of the dataset | Genus level |  | Community level |
|  | *Latilactobacillus* | *Pseudomonas* |  |
| F3M functional genes | 653 | 1087 | 1450 |
| HumanN 3.0 Uniref90 gene families | 409 | 1568 | 1752 |
| F3M metabolic modules | 112 | 156 | 172 |
| HumanN 3.0 metabolic pathways | 30 | 136 | 142 |

\*Values are averages of the four replicates.
\*\*compared to the total number of reads mapping to the whole dataset's *de novo* assembled genes catalog (see materials and methods section).
§ compared to the total number of reads mapping to *L. sakei* or *P. fragi de novo* assembled genes.

Nearly 60% of the total reads from the dataset mapped to the F3M functional annotations, whereas this percentage was 70-80% for the HumanN 3.0 Uniref90 GeneFamilies. This result is not surprising as HumanN 3.0 covers nearly 2,895 Uniref90 GeneFamilies with EC numbers compared to 1,985 functional genes in F3M. However, striking differences are observed when comparing the results obtained on the two main genera. The F3M method gives similar mapping of both genus (50 to 60%) of total genus-specific reads, unlike HUMAnN 3.0 method, which shows reduced efficiency for mapping *Latilactobacillus*-specific reads (∼30%) compared to *Pseudomonas*-specific reads (70-90%). Similarly, the HUMAnN 3.0 method shows important discrepancies in detecting functional categories (gene or pathway level) between the two genera compared to the F3M method. This result unambiguously indicates that the MetaCyc database used as a reference in HumanN 3.0 is biased, as it selects results for certain types of micro-organisms whose metabolic functions have been selected preferentially in the database. Another difference we observed is the considerable difference in the number of MetaCyc metabolic pathways detected by HumanN 3.0 between the two genera and the low number of Uniref90 GeneFamilies efficiently transferred to MetaCyc pathways (only 5-10% of the reads mapping the gene families, data not shown), indicating that many Uniref90 GeneFamilies are not represented in the MetaCyc pathways. In comparison, 100% of reads mapping the F3M functional genes are also mapping the F3M metabolic modules.

To further strengthen the value of the F3M tool suite for food microbiome analysis, we performed a gene expression differential analysis between the two storage conditions using DESeq2. The results obtained at the different levels of data aggregation (Table 4) show that the number of genes/modules over-expressed in vacuum storage condition is higher than in air condition.

**Table 4.** Number of entries from the F3M or HumanN 3.0 functional categories with significant (*p*_*adj*_ < 0.05) differential over-expression between air (log_2_ fold-change < -0.5) and vacuum (log_2_ fold-change > +0.5) in the beef carpaccio dataset.

|  |  | Genus level |  | Community (meta) level |  |
| --- | --- | --- | --- | --- | --- |
|  |  | modules | genes | modules | genes |
| F3M Metabolism Redox & Uptake | AIR | 11 | 48 | 5 | 39 |
|  | VACUUM | 11 | 81 | 10 | 68 |
| F3M House-Keeping | AIR | 8 | 42 | 3 | 30 |
|  | VACUUM | 0 | 44 | 2 | 20 |
| F3M total |  | 30 | 216 | 20 | 157 |
|  |  | pathways | genes | pathways | genes |
| HumanN 3.0 | AIR | 3 | 41 | 6 | 24 |
|  | VACUUM | 3 | 78 | 9 | 73 |
| HumanN 3.0 total |  | 6 | 119 | 15 | 97 |

Finally, by analysing the types of F3M genes, functions or modules differentially expressed, we were able to draw a hypothesis to explain the ecological process that allows *P. fragi* to remain competitive under vacuum conditions (see Figure 6). In summary, *L. sakei* over-expresses catabolic metabolic functions under air storage conditions more than does *P. fragi*. Specifically, this lactic acid bacterium increases the catabolism of carbohydrates (ribose, fructose, and glycerol). Also, it shows an increase in acetate production (significant on the module scale) by several oxidative pathways, such as pyruvate oxidase, pyruvate dehydrogenase, and lactate oxidase. *P. Fragi*, on the other hand, over-expresses catabolism of C4-carboxylic acids and oxidative degradation of fatty acids, indicating separate energy production pathways for the two species and perhaps a low level of competition for energy sources. On the other hand, both *L. sakei* and *P. fragi* species make a metabolic switch towards proteolysis (peptidases and proteases) and amino acid catabolism under vacuum conditions. However, the catabolic repertoire of *P. fragi* for these functional modules is much greater (arginine, lysine, alanine, branch-chain amino acids, and creatine catabolism) than that of *L. sakei*, which may give *P. fragi* a competitive advantage despite the lack of oxygen.

**Figure 6.**
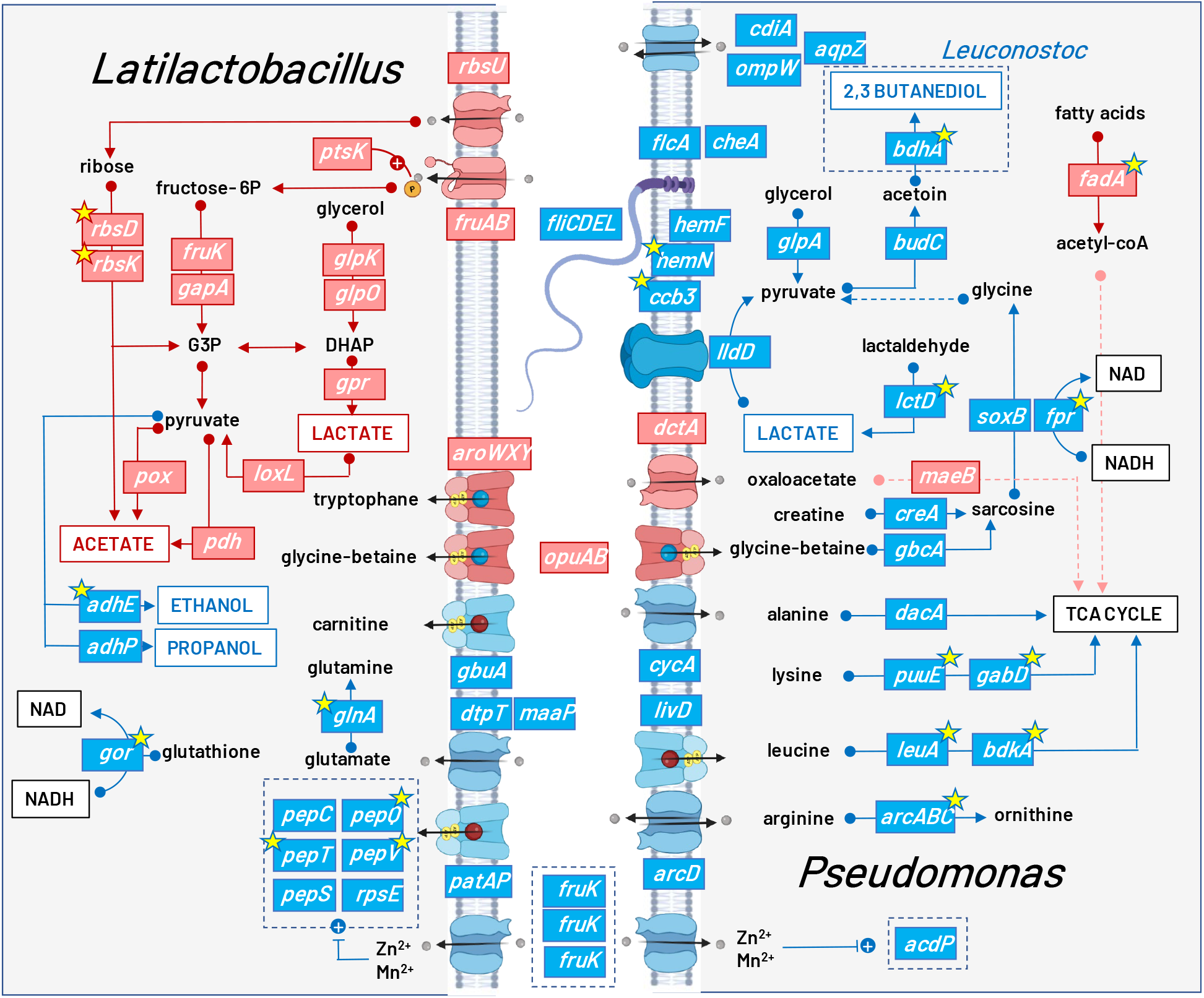
Overview of the metabolic interactions between *L. sakei* (left) and *P. fragi* (right) under air or vacuum storage conditions in beef carpaccio deciphered by the F3M tool suite. The figure illustrates around 60 metabolic functions selectively chosen to highlight those overexpressed in air (depicted by red boxes) and those overexpressed in vacuum (depicted in blue boxes). Uptake systems are shown for both species along the cell membrane, illustrated in the center. The yellow stars indicate the functional genes that were also identified by the HUMAnN 3.0 method and subsequent DESeq2 analysis.

The analysis also showed that, under vacuum conditions, *P. fragi* increased the expression of its main respiratory chain (ccb3-type, *i*.*e*., O_2_-dependent) and its heme co-factor (*hemN/F*) in order to use lactate as an energy and carbon source. As lactate is one of the main fermentation products of *L. sakei*, this metabolic exchange is beneficial to *P. fragi*. Indeed, the trophic chain may be extended downstream to *Leuconostoc gelidum* (a sub-dominant species detected in this simplified ecosystem) through the subsequent production of 2,3-butanediol from acetoin, itself produced by *P. fragi* from pyruvate. However, using the oxygen-dependent cbb3-type respiratory chain under vacuum conditions remains a surprising observation. In our opinion, this result might suggest the existence of a residual level of oxygen in vacuum-packed meat (particularly rich in heme, on which oxygen may remain chelated), but that this concentration of O_2_, probably sub-optimal, induces over-expression of chemotaxis and flagellum synthesis in *P. fragi* (*che* and *fli* functions) for efficient recovery.

Finally, the results of this dataset reveal the added value of the uptake and redox modules specific to the F3M tool suite. On the one hand, the differentially expressed transport systems are perfectly consistent with the metabolic pathways activated in each species, and in some cases, additional additional features of the metabolic network regulation were revealed, such as the activation of zinc transporters in *L. sakei* and *P. fragi* under vacuum conditions. Zinc is one of the co-factors most frequently used by peptidases and proteases. The redox module also indicates that, under vacuum conditions, the two bacteria activate two different systems to help re-oxidise the NADH produced by catabolic pathways: the reduction of glutathione disulfide (abundant in meat) by *L. sakei* and the flavodoxin reductase pathway (linked to creatine catabolism) by *P. fragi*.

## Discussion and conclusions

The analysis of metabolic interactions within food microbiomes poses a number of challenges, including the enormous diversity of identifiable microbial communities between samples, the significant variability in abundance between species, and, above all, the difficulty of automatically inferring the annotation of certain anaerobic fermentation-type metabolic functions based on available generalist databases such as KEGG or MetaCyc.

The F3M tool suite aims to fill this gap and to build an intuitive, easy-to-use, open-access tool that will make it easy to aggregate gene expression information at different functional and taxonomic levels. Therefore, the F3M tool suite was designed to be modular in terms of the choice of gene catalogs to be inferred and the level of aggregation of expression data. Analysis of the dataset produced from the microbiomes of beef carpaccio stored under two conditions confirms that using F3M quickly generates information that can be interpreted in terms of hypotheses concerning metabolic interactions between species.

As with other tools based on the same principle of functional modules (Shiroma et al., 2024; Valles-Colomer et al., 2019; Vieira-Silva et al., 2016), the added value of F3M comes from the extensive manual curation of the key functions of food microbiomes and their association into modules that facilitate the interpretation of interactions. We have chosen to limit the number of genes and functions curated in this first version of the database (1,427 and 865, respectively, if we exclude the housekeeping modules, a limited number compared with the general metabolic databases available). This choice was guided, on the one hand, by the need to verify the curation process and, on the other hand, in order to achieve targeted filtering of the key functions that predominate in food microbiomes. This process may also represent a limitation, as many metabolic functions remain to be included in the database. For example, we chose not to annotate multidrug transporters and efflux pumps. The low degree of precision of the annotations and the enormous degree of paralogy within this functional class made curation impractical at this time. We also chose to limit the expertise of CAZYmes by selecting only those enzymes which were most relevant in the context of food microbiomes. More in-depth expertise on this functional category can nevertheless be carried out in parallel with the appropriate authoritative CAZy database and cognate tools (Drula et al., 2022; Hobbs et al., 2023; Lombard et al., 2025, Lombard et al., 2014), and we plan to improve their inclusion in the F3M in further updates.

As mentioned above, in keeping with the great variability of food ecosystems, we were also confronted with metabolic pathways not covered by the automatic annotation of the eggNOG mapper. For example, we identified in some *Lactobacillaceae* genomes the pathway for lysine catabolism through the saccharopine pathway, which degrades lysine to alpha-aminoadipate (Serrano et al., 2012). The enzymes of this pathway are identified in the KEGG database and are associated with KO numbers, but are not present in MetaCyc. The information in KEGG, however, refers only to this pathway as it pertains to humans and plants. Although Serrano and co-authors described the existence of this pathway in bacteria, the bacterial orthologous genes were not identified using the eggNOG mapper tool due to a lack of bacterial genes correctly annotated for this pathway. Hence, in this and similar cases, curation can only be considered once KO identifiers have been established in the generic databases.

The dependence of the F3M strategy on eggNOG mapper is an important element to consider for the evolution of our tools. Undoubtedly, the eggNOG mapper annotation process will improve as the scientific community updates the annotations in KEGG and MetaCyc. Nevertheless, if this tool changes too significantly in its structure and annotation workflow, it will be necessary to update our database codification and builder.

We aim to proceed stepwise and to make progressive improvements to the F3M tool suite with frequent updates. One of the first steps will be to include genes and metabolic functions carried by yeasts and other fungi, as the F3M database is currently exclusively bacterial. This improvement step represents a significant challenge, as the functional annotation of genes from eukaryotic micro-organisms requires more in-depth expertise than their bacterial counterparts. However, this scientific field is changing rapidly, both because of the greater effort being made to characterise the genomic diversity of yeasts and fungi in food microbiomes and because of the development of suitable tools for detecting eukaryotic genes in omics datasets. It is, therefore, possible to anticipate that as the process of annotating yeast and fungal genes using eggNOG mapper and other tools improves, this will allow improving the builder framework to implement more diverse gene catalogs or sources of genomic data.

The interoperability of the F3M tools suite functional annotations with other references, such as KEGG and Pathways tools, remains an important element in the evolution process of F3M. We plan to rapidly integrate cross-references (metabolite codes and numbers, for instance) and continue to rely on existing databases to offer, in return, our annotation expertise. In this respect, we also intend to build on initiatives such as the Fermentation Explorer (Hackmann and Zhan, 2023), whose objectives are more focused on annotation inference for anaerobic metabolisms.

Another critical step in the evolution of the F3M modules will be to implement the knowledge from RNA-seq datasets. Learning from food microbiome gene expression experiments will be one of the most effective strategies for resolving the limitations currently associated with the selected and curated modules, notably by improving annotation. The use, in parallel, of other software such as scoary2 (Roder et al., 2024), which enables the connectivity and integration of phenotypic and metabolomic datasets using ultra-fast microbial genome-wide association studies, is certainly a strategy worth considering. We also believe the F3M tool suite will provide key information for improving genome-scale metabolic modeling strategies applied to food microbiomes. Similarly, a benchmark analysis of various RNA-seq datasets, comparing F3M and other tools, will enable us to better define the advantages and limitations of F3M to improve its structure and content.

## Acknowledgements

We are grateful to the INRAE MIGALE bioinformatics facility (MIGALE, INRAE, 2020. Migale Bioinformatics Facility, doi:10.15454/1.5572390655343293E12) for providing help and/or computing and/or storage resources.

## Data, scripts, code, and supplementary information availability

The F3M database, the gene catalogs and the tutorial dataset are licensed under Etalab 2.0; the F3M builder tool and f3mr R package are licensed under MIT.

The F3M database and the supplementary file containing the metabolic maps of the various modules are available at the Food Microbial Ecology lab dataverse: https://doi.org/10.57745/9VKS65.

Scripts, code, and documentation (Wiki and troubleshooting) for the F3M builder are available at: https://forge.inrae.fr/fme_team/f3m_builder.

The f3mr R package and relevant documentation are available at: https://forge.inrae.fr/fme_team/f3mr and at the Food Microbial Ecology lab website: https://fme_team.pages-forge.inrae.fr/f3mr/.

The two F3M-inferred gene catalogs dedicated to studies on meat microbiomes and fermented vegetables microbiomes are available at the Food Microbial Ecology lab dataverse: https://doi.org/10.57745/7T05VI and https://doi.org/10.57745/IOADXW, respectively.

The tutorial beef carpaccio food metatranscriptomic dataset and the corresponding metadata are available at the Food Microbial Ecology lab dataverse: https://doi.org/10.57745/STFX9L.

